# *De novo* Transcriptome Characterization of *Iris atropurpurea* (the Royal Iris, *Iris* section *Oncocyclus*) and Phylogenetic Analysis of MADS-box and R2R3-MYB Gene Families

**DOI:** 10.1101/680363

**Authors:** Bar-Lev Yamit, Senden Esther, Pasmanik-Chor Metsada, Sapir Yuval

## Abstract

The Royal Irises, *Iris* section *Oncocyclus*, are a Middle-Eastern group of irises, characterized by extremely large flowers with a huge range of flower colors and a unique pollination system. The Royal Irises are considered to be in the course of speciation and serve as a model for evolutionary processes of speciation and pollination ecology. However, no transcriptomic and genomic data for molecular characterization are available for these plants.

Transcriptome sequencing is a valuable resource for determining the genetic basis of ecological-meaningful traits, especially in non-model organisms. Here we describe the *de novo* transcriptome sequencing and assembly of *Iris atropurpurea*, an endangered species, endemic to Israel’s coastal plain. We employed RNA sequencing to analyze the transcriptomes of roots, leaves, and three stages of developing flower buds. To identify genes involved in developmental processes we generated phylogenetic gene trees for two major gene families, the MADS-box and MYB transcription factors, which play an important role in plant development. In addition, we identified 1,503 short sequence repeats that can be developed for molecular markers for population genetics in irises.

In the era of large genetic datasets, the *Iris* transcriptome sequencing provides a valuable resource for studying adaptation-associated traits in this non-model plant. This first reported transcriptome for the Royal Irises, and the data generated from this study, will facilitate gene discovery, functional genomic studies, and development of molecular markers in irises, to complete the intensive eco-evolutionary studies of this group.

## Introduction

*Iris* is the largest genus in the Iridaceae (Asparagales) with over 300 species (Makarevitch, Golovnina, Scherbik, & Blinov, 2003; Matthews, 1997). The genus is highly heterogeneous, with species exhibiting a wide range of plant sizes, and flower shapes and colors (Matthews, 1997).

The Royal Irises (*Iris* section *Oncocyclus*) are a Middle-Eastern group of about 32 species that are endemics to dry, Mediterranean-type climates and found in the eastern Mediterranean Basin, Caucasica, and central Anatolia (Carol A. Wilson, Padiernos, & Sapir, 2016). Species of section *Oncocyclus* in Israel occur in small isolated populations and many are considered rare, threatened, or endangered (Shmida & Pollak, 2007). These species are characterized by a single large flower on a stem and perennial, short, knobby rhizomes, occasionally with stolons (Sapir & Shmida, 2002; Carol A. Wilson et al., 2016). Plants are diploid with chromosome number of 2n=20 (Avishai & Zohary, 1977). This number is relatively low for *Iris* species, whose chromosome number ranges from 2n = 16 in *I. attica* to 2n = 108 in *I. versicolor* (data obtained from Chromosome Count DataBase (Rice et al., 2015)), and genome size ranges from 2,000 to 30,000 Mbp (Kentner, Arnold, & Wessler, 2003).

The Royal Irises are thought to be undergoing recent speciation (Avishai & Zohary, 1980; Sapir & Shmida, 2002; Carol A. Wilson et al., 2016). Consequently, in recent years, they have emerged as a platform for the study of evolutionary processes of speciation, adaptation and pollination ecology (Arafeh et al., 2002; Dorman, Sapir, & Volis, 2009; Lavi & Sapir, 2015; Sapir & Mazzucco, 2012; Sapir, Shmida, & Ne’eman, 2005, 2006; Volis, Blecher, & Sapir, 2010; Carol A. Wilson et al., 2016; Yardeni, Tessler, Imbert, & Sapir, 2016). Evolutionary processes and adaptive phenotypes are governed by genetic differences. Thus, the study of plant ecology and evolution increasingly depends on molecular approaches, from identifying the genes underlying adaptation, reproductive isolation, and speciation, to population genetics. No genetic and molecular tools are yet available for the Royal Irises. Whole-genome sequencing of the *Iris* is a challenging task, due to its large genome size (Kentner et al., 2003), and therefore transcriptome sequencing may provide a feasible, still a strong genomic resource.

Transcriptome sequencing is a powerful tool for high-throughput gene discovery, and for uncovering the molecular basis of biological functions, in non-model organisms (Jain, 2011). Few *Iris* transcriptomes have been already sequenced (Ballerini, Mockaitis, & Arnold, 2013; S. Tian et al., 2015) (C.-S. Gu et al., 2017; C. Gu et al., 2018), all are of irises which are in distant clades from *Oncocyclus* iris (Carol A. Wilson, 2011). Currently, only one NGS-based dataset is available for the Royal Irises, which is a plastid genome sequence of *Iris gatesii* (Carol A Wilson, 2014). Previous attempts to transfer molecular tools developed for Louisiana irises to *Oncocyclus* irises, such as the development of microsatellite loci or identifying candidate genes, have failed (Y. Sapir, un-published). Furthermore, Royal Iris species have low plastid variance (Y. Sapir and Y. Bar-Lev, un-published) and lack nuclear sequences. All these, call for a wider set of molecular tools. Our main objective was to generate a reference RNA sequence for the Royal Irises that can serve as a molecular toolbox.

Here we report the *de novo* assembly of a transcriptome for *Iris atropurpurea* Baker, one of the Royal Irises species. *I. atropurpurea* is a highly endangered plant endemic to Israeli coastal plain (Sapir, 2016; Sapir, Shmida, & Fragman, 2003). In recent years this species has been studied extensively for its morphology (Sapir & Shmida, 2002), pollination (Sapir et al., 2005, 2006; Watts, Sapir, Segal, & Dafni, 2013), speciation, and population divergence (Sapir & Mazzucco, 2012; Yardeni et al., 2016). In order to answer any further questions in this system, molecular tools are needed. Transcriptome sequencing of *I. atropurpurea* will facilitate further studies of genetic rescue, population genetics, as well as finding genes that underlie different biological functions.

One of the most important biological functions to understand plant evolution is plant development. To identify genes involved in developmental processes, we analyzed the phylogeny of sequences annotated to MADS-box and R2R3-MYB transcription factors families, which are involved in the regulation of diverse developmental functions. Homologs for genes of these families have been identified in *Iris fulva* of the Louisiana irises (Ballerini et al., 2013). We therefore aimed to identify their homologs in the Royal Irises.

Plant development greatly depends on the function of MADS-box transcription factors, a very ancient family of DNA binding proteins, which are present in nearly all major eukaryotic groups. MADS-box genes comprise a highly conserved sequence of ~ 180 bp, which encodes the DNA binding domain in the MADS-box protein (Glover, 2014; Heijmans, Morel, & Vandenbussche, 2012). MADS-box genes are divided into type I and type II. In plants, type I MADS-box genes are subdivided into three groups: Mα, Mβ and Mγ (De Bodt, Raes, Florquin, et al., 2003; Par◻enicová et al., 2003). They are involved in female gametophyte, embryo sac, and seed development. The type II MADS-box genes in plants are known as the MIKC MADS-box group and are extensively studied. MIKC proteins convey three additional distinctive regions: an intervening region (I), a keratin-like domain (K), and a C-terminal domain (C) (Gramzow, Ritz, & Theißen, 2010; Gramzow & Theissen, 2010). Found within this group are the MIKCc and MIKC* subgroups (Henschel et al., 2002). MIKCc MADS-box genes (the c stands for classic), are mainly involved in plant and flower development (Coen & Meyerowitz, 1991; Schwarz-Sommer, Huijser, Nacken, Saedler, & Sommer, 1990), and are phylogenetically divided into 14 major groups in *Arabidopsis* and rice(Arora et al., 2007; Becker & Theißen, 2003). The MIKC* group, in some reports, matches the *Arabidopsis* Mδ subgroup, defined as part of the type I group (De Bodt, Raes, Van de Peer, & Theißen, 2003).

Another superfamily of transcription factors, that are important for plant development, are the MYB proteins, which contain the conserved MYB DNA-binding domain (Stracke, Werber, & Weisshaar, 2001). The MYB family members are categorized based on the number of MYB domain repeats: 1R-(MYB related genes, containing a single or partial MYB domain), R2R3-, 3R- and 4R-MYB proteins (Dubos et al., 2010; Stracke et al., 2001; Yanhui et al., 2006). MYB proteins are widely distributed in plants, in which the R2R3-MYB subfamily is the most abundant (containing an R2 and R3 MYB domain) (Ambawat, Sharma, Yadav, & Yadav, 2013; Dubos et al., 2010; Stracke et al., 2001). The large abundance of the R2R3-MYB family in plants indicates their importance in the control of various plant specific processes, such as responses to biotic and abiotic stresses, development, defense reactions, flavonoid and anthocyanin biosynthesis, regulation of meristem formation, and floral and seed development (reviewed in(Ambawat et al., 2013) and (Du et al., 2009)).

Here we employed phylogenetic approach to identify homologs of MADS-box and R2R3-MYB transcription factors in the *I. atropurpurea* transcripts. We sequenced transcriptomes from various tissues and flower bud developmental stages. From these, we established an annotated database for *I. atropurpurea*, potentially applicable to other species of the Royal Irises, and explored the homologs of MADS-box and R2R3-MYB. This is the first reported transcriptomes for the *Oncocyclus* section. The sequenced *Iris* transcriptome offers a new foundation for genetic studies and enables exploring new research questions.

## Methods

### Plant material

We used two accessions (genotypes) of *I. atropurpurea*, DR14 and DR8. Plants were brought from a large *I. atropurpurea* population in Dora (32°17’N 34°50’E) in Israel (figure 1a) and grown at the Tel Aviv University Botanical Garden. Aiming at finding genes related to flower development and floral traits, we used three different bud developmental stages. We defined bud developmental stage 1 as the earliest detectable bud, where the bud has no color. Stage 2 is a bud around 1.5 cm in size with the anthers still prominently visible above the petals, and at the onset of color production. Stage 3 is a full-colored bud, over 2 cm in size and with the petals covering the anthers (figure 1b). Earlier stages of flower development in the Royal Irises are nearly impossible to detect in naturally-growing plants. In these stages the meristem is attached to the rhizome underground and requires much destruction of the plant to be found (Perl, 1984). We collected tissues from the root, young leaf and four buds in three developmental stages (one bud from stages 1 and 3, and two buds of stage 2) from DR14. We also collected buds in stages 1 and 2 from DR8 to enlarge the representation of rare or low expressed genes. Unfortunately, due to the low number of flowers (buds) per plant in Royal Irises, we were unable to obtain replicates for all bud stages.

**Figure 1.**
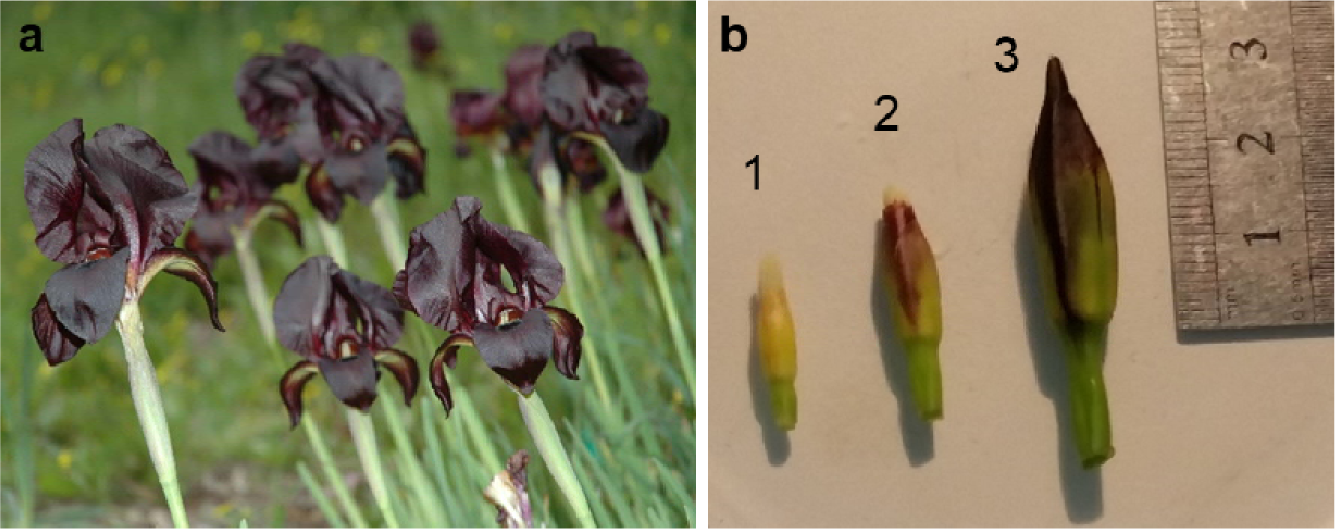
Plant materials used for RNA sequencing. a. *Iris atropurpurea* flower in the field site where collected (Dora). b. Representation of three stages of bud development (1 to 3) in *I. atropurpurea*, as defined in the text.

### RNA isolation and sequencing

We extracted total RNA from all the tissue samples using the RNeasy Mini Kit (Qiagen, Hilden, Germany), according to the manufacturer’s instructions. We measured the quantity and quality of each RNA sample using Qubit fluorometer (Invitrogen) and Bioanalyzer TapeStation 2200 (Agilent Technologies Inc., USA), respectively. Only RNA samples that presented sufficient 260/280 and 260/230 purity and RIN (RNA integrity number) above 8.0 were used for sequencing. RNA was processed by the Technion Genome Center as following: RNA libraries were prepared using TruSeq RNA Library Prep Kit v2 (Illumina), according to manufacturer’s instructions, and libraries were sequenced using HiSeq 2500 (Illumina) on one lane of 100 PE run, using HiSeq V4 reagents (Illumina). Sequences generated in this study were deposited in NCBI’s Gene Expression Omnibus (GEO, http://www.ncbi.nlm.nih.gov/geo/) under the GEO accession number GSE121786.

### *De novo* transcriptome assembly and annotation

The quality of the raw sequence reads was estimated using FastQC. De novo assembly of the *Iris* transcriptome was done using Trinity (version trinityrnaseq_r20140717), with a minimum contig length of 200 base pairs (bp) (Grabherr et al., 2011). We estimated assembly quality using Quast (v. 3.2) (Gurevich, Saveliev, Vyahhi, & Tesler, 2013). Contigs (isoforms) that are likely to be derived from alternative splice forms or closely-related paralogs were clustered together by Trinity and referred to as “transcripts”. The initial reads from each sample were mapped back to the *Iris* transcriptome that was assembled, using trinity pipeline and Bowtie (v. 1.0.0). The number of mapped reads per transcript per sample was counted using RSEM (v. 1.2.25) (B. Li & Dewey, 2011).

To find the putative genes and function, transcripts were aligned against the UniProt non-redundant protein database (26-09-2016) and against PFAM protein family database (El-Gebali et al., 2018), using BLASTX alignment with an e-value cutoff to < 0.0001 (Altschul, Gish, Miller, Myers, & Lipman, 1990). To classify functions of the transcripts, they were also aligned against Clusters of Orthologous Groups (COGs) protein database (ftp://ftp.ncbi.nih.gov/pub/COG/COG/). For transcription factors prediction, the we submitted the predicted protein sequences to search against PlantTFDB (http://planttfdb.cbi.pku.edu.cn/index.php) (F. Tian, Yang, Meng, Jin, & Gao, 2019).

### Phylogeny analysis

We retrieved all *Iris* transcripts that were annotated as either MADS or MYB proteins from the transcriptome and translated the longest open reading frame (ORF) using Virtual Ribosome (Wernersson, 2006). We took the transcriptome transcripts that also contain the MADS or R2R3-MYB domain by PFAM. We downloaded *I. fulva* protein sequences for MIKCc MADS-box and R2R3-MYB transcription factors from NCBI (Ballerini et al., 2013). *Arabidopsis* and rice (*oryza sativa*) MADS and R2R3-MYB sequences were taken from their genome databases [The *Arabidopsis* Information Resource (TAIR): www.arabidopsis.org and the Rice Genome Annotation Project (RGAP): rice.plantbiology.msu.edu, respectively]. The gene identifiers were denoted to AtMYB genes in *Arabidopsis* and the locus id in rice to avoid confusion when multiple names are used for same gene. The sequences of each gene family were trimmed using trimAl v1.3 (Capella-Gutiérrez, Silla-Martínez, & Gabaldón, 2009) and aligned using ClustalW alignment (Thompson, Higgins, & Gibson, 1994), in MEGA X Molecular Evolutionary Genetics Analysis Software (Kumar, Stecher, Li, Knyaz, & Tamura, 2018). We tested for the best substitution model and found that the best model for MADS is the JTT (Jones, Taylor, Thornton) model (Jones, Taylor, & Thornton, 1992) + Gamma-distributed rates (G), and for MYB, JTT + G + amino acid frequency (F). For comparative phylogenetic analysis, we used maximum likelihood in MEGA X (Kumar et al., 2018) with 1000 bootstrap replications. Phylogenetic trees were visualized using FigTree v1.4.3 (Rambaut, 2007).

### SSRs mining

In order to utilize the transcriptome sequenced also for population genetic markers, we searched for short sequence repeats (SSRs; microsatellites) in the assembled contigs. We used a Perl script (find_ssrs.pl; (Barker et al., 2010)) to identify microsatellites in the unigenes. In this study, SSRs were considered to contain motifs with two to six nucleotides in size and a minimum of four contiguous repeat units.

## Results and Discussion

### Sequencing of *Iris* transcriptome

To generate the *Iris* transcriptome, eight cDNA libraries were sequenced: root, leaf and three bud stages from one genotype of *I. atropurpurea* (DR14), and buds in stages 1 and 2 from a different genotype of the same population (DR8). We generated a total of 195,412,179 sequence reads. The average GC content of *Iris* contigs was 47% (table 1 and 2). Reads were of very high quality throughout their length, without evidence of adapter content (Phred score >30).

**Table 1.** Statistical summary of *Iris* transcriptome sequencing and assembly.

|  |  |
| --- | --- |
| Total reads | 195,412,179 |
| Contigs (Isoforms) | 258,466 |
| Transcripts | 184,341 |
| Transcriptome size | 168,049,166 |
| N50 contig size ($\geq 500$ bp) | 1,312 |
| Largest contig | 27,971 |

**Table 2.** Descriptive statistics of *Iris* transcriptome samples. GC – Percentage of G or C nucleotides in the sequence.

| Plant ID | Tissue | # Paired-end sequences | #Reads | %GC | Total mapped reads | % Unique mapped reads |
| --- | --- | --- | --- | --- | --- | --- |
| **DR14** | Root | 47,760,556 | 23,880,278 | 48 | 16,671,086 | 55 |
|  | Leaf | 52,936,482 | 26,468,241 | 47 | 18,366,659 | 54 |
|  | Bud stage 1 | 54,806,560 | 27,403,280 | 47 | 16,610,741 | 40 |
|  | Bud stage 2 (a) | 51,092,390 | 25,546,195 | 47 | 16,095,948 | 43 |
|  | Bud stage 2 (b) | 48,073,466 | 24,036,733 | 47 | 15,363,667 | 43 |
|  | Bud stage 3 | 59,105,996 | 29,552,998 | 47 | 18,570,119 | 43 |
| **DR8** | Bud stage 1 | 36,949,342 | 18,474,671 | 45 | 10,887,290 | 40 |
|  | Bud stage 2 | 40,099,566 | 20,049,783 | 46 | 12,509,415 | 45 |

Using Trinity, we assembled 258,466 contigs (isoforms) longer than 200 bp, which clustered into 184,341 transcripts, with a total length of 168,049,166 bp. A larger N50 length and average length are considered indicative of better assembly. The longest contig was 27,971 bp and half of the contigs (N50) with more than 500 bp were above 1,312 bp long (table 1).

To quantify the abundance of contigs assembled, the reads of the separated *Iris* organs were mapped to the assembled contigs, with 125,074,925 mapped reads overall, and an average of 45% reads per tissue that mapped to a unique sequence in the assembled transcriptome.

The length distribution of the assembled contigs revealed that 126,194 (68.46%) contigs ranged from 201 to 500 bp in length; 37,335 (20.25%) contigs ranged from 501 to 1,000 bp in length; 16,282 (8.83%) contigs ranged from 1,001 to 2,000 bp in length; and 4,530 (2.46%) contigs were more than 2,000 bp in length (figure 2). Descriptive statistics of the sequencing data and transcriptome assembly are summarized in tables 1 and 2.

**Figure 2.**
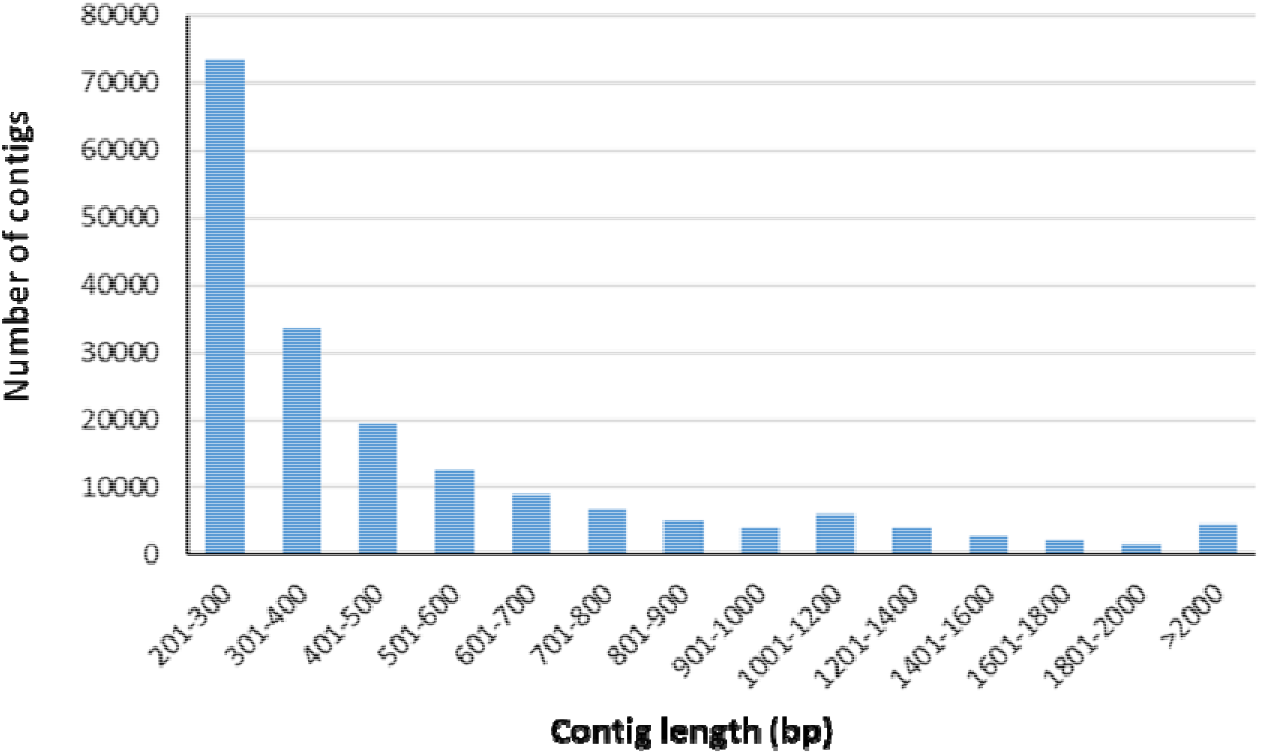
Distribution of contig lengths (in base pairs) across the assembled contigs from the *Iris* transcriptome.

### Annotation of *Iris* transcriptome

Using BLASTX search against the UniProt database, we identified 28,708 transcripts with at least one significant hit. Transcripts mostly annotated to *Arabidopsis thaliana* (%), *Oryza sativa* Japonica Group (%) and *Nicotiana tabacum* (%) (figure 3a). Surprisingly, a significant proportion of the annotated transcripts were annotated as *Arabidopsis thaliana*, while only 10 % were annotated as *Oryza sativa*, which is a monocot and therefore more closely related to irises. This is probably due to the higher representation of genomic resources for *Arabidopsis thaliana*. A considerable number of transcripts annotated to “non-plant” organisms, most of them to human (*Homo sapiens*, 2. %) (figure 3a). This may be attributed to housekeeping genes, which are preserved across all species in eukaryotes, and may also be due to the highly annotated human genome.

**Figure 3.**
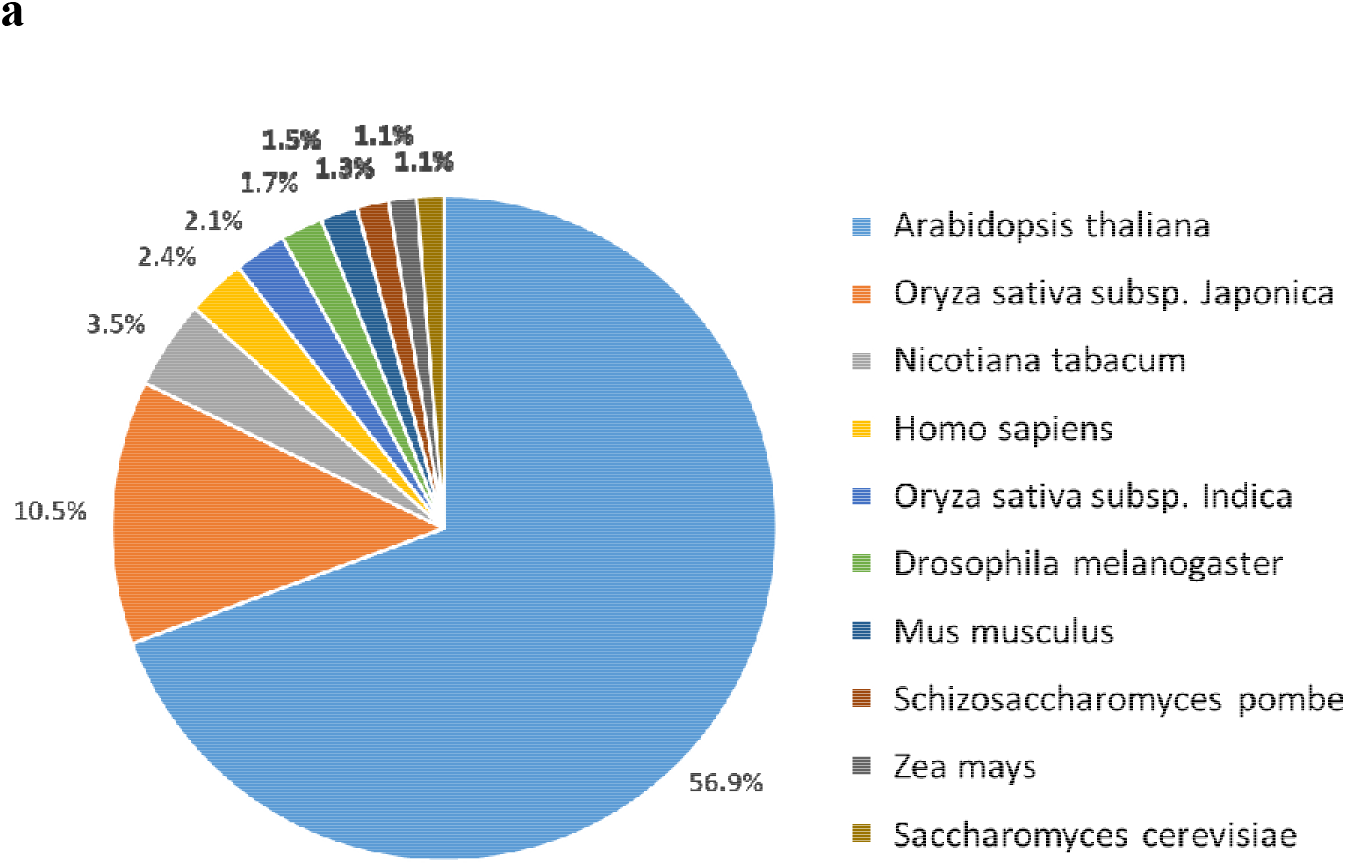
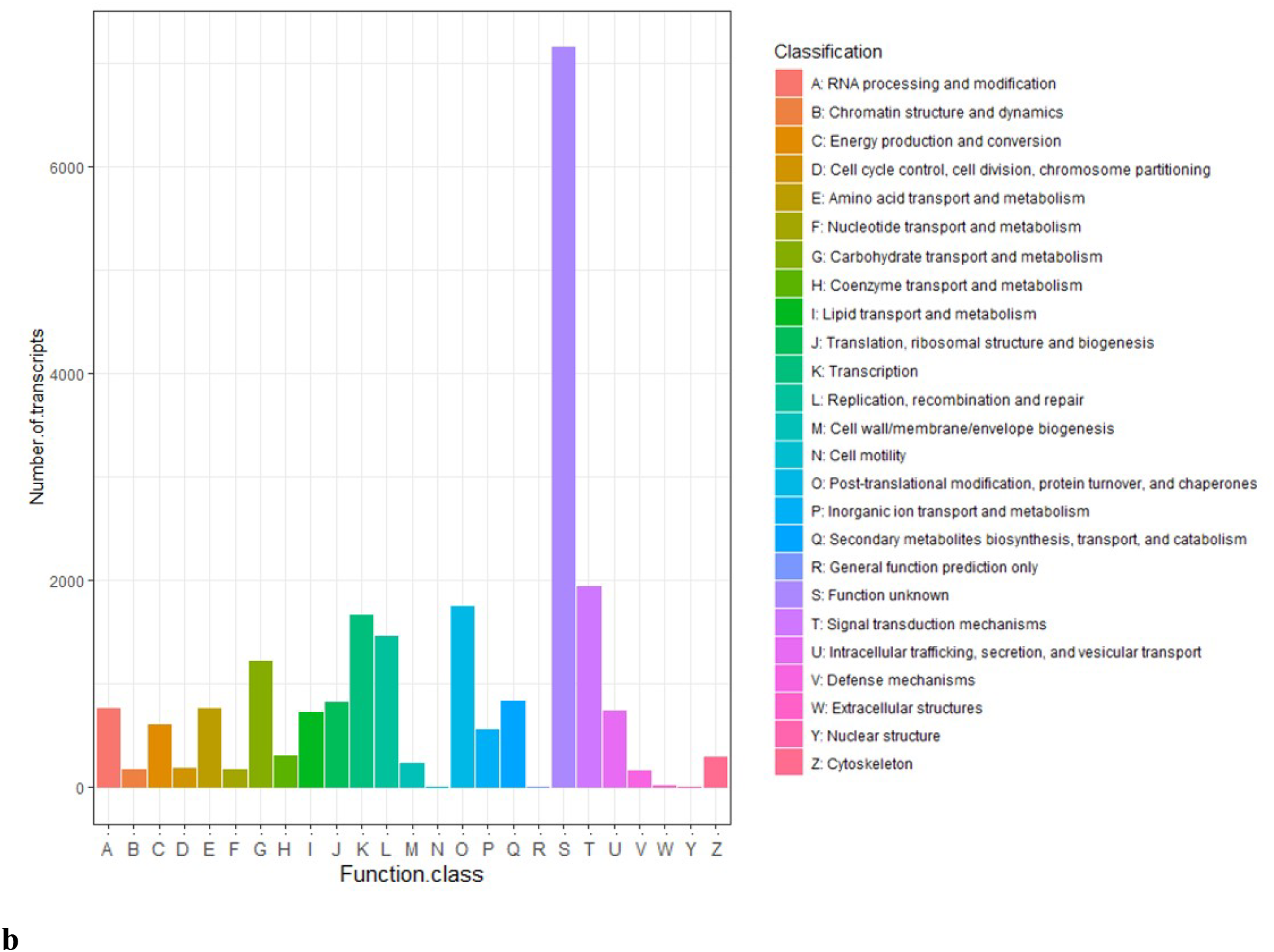
a. Top 10-hit species distribution of annotated transcripts. Other species represented in the transcriptome had only 1% or less of the transcripts annotated to them. b. Clusters of orthologous group (COG) classification, showing 22,564 transcripts that were classified.

Search against the COG database resulted in the classification of 22,564 transcripts (table S1). Among the 25 COG categories, the cluster for unknown function was the largest group (7,151, 31.69%). The following categories of the top ten are: signal transduction mechanisms (1938, 8.59%), posttranslational modification, protein turnover and chaperones (1754, 7.77%), transcription (1670, 7.4%), replication, recombination and repair (1461, 6.47%), carbohydrate transport and metabolism (1222, 5.42%), secondary metabolites biosynthesis, transport and catabolism (839, 3.72%), translation, ribosomal structure and biogenesis (825, 3.66%), amino acid transport and metabolism (767, 3.4%), and RNA processing and modification (764, 3.39%) (figure 3b). The COG term ‘signal transduction’ was also enriched in previous transcriptomes, such as in *Iris lactea* (C. Gu et al., 2018), *Camelina sativa* L (Mudalkar, Golla, Ghatty, & Reddy, 2014), and in *Taxodium* ‘Zhongshanshan 405’ (Yu, Xu, & Yin, 2016).

In the PFAM analysis, we found 17,385 (9.43%) *Iris* transcripts that contain at least one PFAM protein domain, and that were classified into 3399 Pfam domains/families (table S1). The 10 most abundant protein families in *I. atropurpurea* are Pkinase, PPR_2, Pkinase_Tyr, LRR_8, RRM_1, RVT_1, PPR_1, p450, PPR3, and LRRNT_2 (figure 4a). Among these protein domains/families, “Protein kinase” and “Tyrosine-protein kinase”, were highly represented. These proteins are known to regulate the activation of most cellular processes (Lehti-Shiu & Shiu, 2012), indicating active signal transduction. This is in accordance with our COG results, also showing enrichment of signal transduction genes. Top ranked family is also PPR_2 -pentatricopeptide repeats. The PPR family controls varied features of RNA metabolism and plays a profound role in organelle biogenesis and function, e.g. mitochondria and chloroplasts (Filipovska & Rackham, 2013; Lurin et al., 2004) (Barkan & Small, 2014). Thus, PPR’s have an essential effect on photosynthesis, respiration, plant development, and environmental responses (Barkan & Small, 2014).

**Figure 4.**
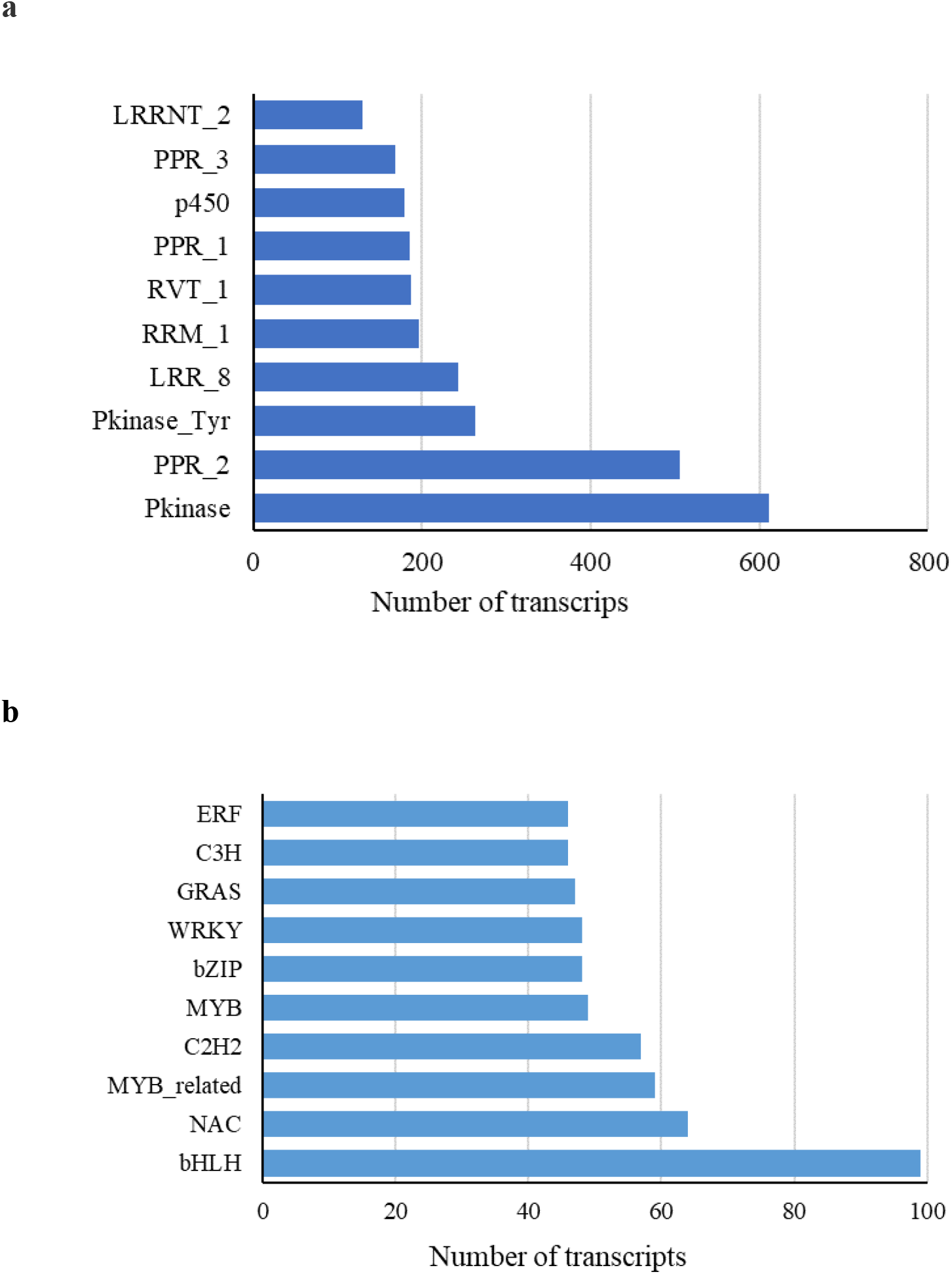
a. The 10 most abundant PFAM protein families in the *I. atropurpurea* a transcriptome. b. The 10 most abundant transcription factors families in the *I. atropurpurea* transcriptome.

Transcription factors (TFs) are key regulators in biological processes. For prediction of transcription factors, we assigned the protein sequences of all the transcripts to PlantTFDB (F. Tian et al., 2019). We found 1021 transcripts that are predicted to be involved in transcription regulation and were classified into 54 transcription factor families (figure 4b, table S1). The basic helix–loop–helix (bHLH) transcription factors family was the most abundant in *I. atropurpurea* consisting of 99 gene family members. In plants, the bHLH proteins are associated with a variety of developmental processes, such as trichomes development (Morohashi et al., 2007; M. Zhao, Morohashi, Hatlestad, Grotewold, & Lloyd, 2008), phytochrome signaling(Duek & Fankhauser, 2005), and cell proliferation and differentiation (Morohashi et al., 2007; Vera-Sirera et al., 2015). bHLH proteins have also been shown to interact with other transcription factors such as MYB (M. Zhao et al., 2008; Zimmermann, Heim, Weisshaar, & Uhrig, 2004). Furthermore, a protein complex of bHLH and MYB transcription factors, associated with a WD40 repeat protein, regulates various cell differentiation pathways and the anthocyanin biosynthesis pathway (Goff, Cone, & Chandler, 1992; Ramsay & Glover, 2005). The rest of the top 10 TFs are: NAC, MYB-related, C2H2, MYB, bZIP, WRKY, GRAS, C3H and ERF.

In total, we identified 33,033 transcripts in at least one database. We were unable to annotate or give a functional prediction to a large fraction of the transcripts. These transcripts could be *Iris* specific genes, genes that have diverged considerably, or genes that are not yet identified in plants.

### Phylogenetic analysis of MADS-box and R2R3-MYB gene families

In the search for orthologous genes involved in flower development in irises, we phylogenetically analyzed two major transcription factor groups, the MADS-box and MYB protein families, to validate the subfamily identities of these genes from *I. atropurpurea*. We performed the phylogenetic analyses using MADS-box and R2R3-MYB protein sequences from *Arabidopsis thaliana* and rice (*Oryza sativa*), the top two annotated species in the transcriptome, and from *Iris fulva*.

### MADS-box genes

MADS-box proteins, and their complex function, regulate floral organ characteristics and are essential for flower development (Heijmans et al., 2012; Honma & Goto, 2001; Theißen & Saedler, 2001). In the *Iris atropurpurea* transcriptome, 43 transcripts were annotated as belonging to the MADS-box family and/or contain the MADS domain. Phylogenetic analysis using *Arabidopsis*, rice, and *I. fulva,* shows orthologous of *I. atropurpurea* in almost all clades of MADS-box proteins (figure 5). The general organization for most clades was similar to previous comparative phylogenies (Arora et al., 2007; Ballerini et al., 2013).

**Figure 5.**
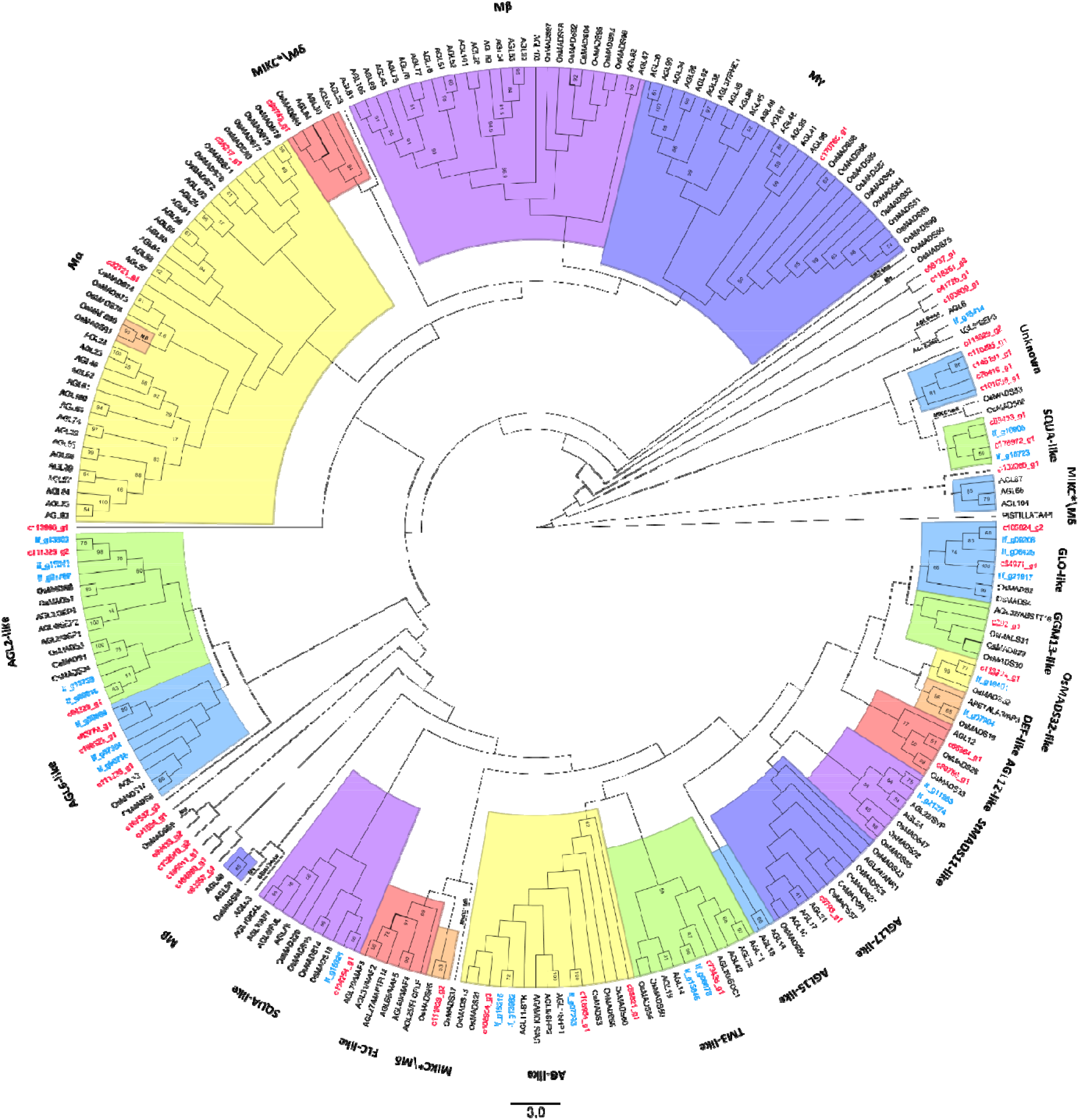
Phylogenetic analysis of MADS-box proteins from the *I. atropurpurea* transcriptome, *I. fulva, Arabidopsis* and rice. *I. atropurpurea* transcripts names are in red and *I. fulva* in light blue. Colours are for visual separation only. Sequences that were separated from their known clade have the name of their original clade written on the branch.

Of the *I. atropurpurea* MADS-box genes identified, 19 clustered with MIKCc, 2 with Mα, 0 with Mβ, 1 with Mγ, and 2 grouped with MIKC*/ Mδ-type genes. Among the genes clustered with type II MIKCc MADS, we identified all 14 documented clades (Arora et al., 2007; Ballerini et al., 2013; Becker & Theißen, 2003)), comprising representative genes of *Arabidopsis*, rice, and *I. fulva*. *I. atropurpurea* had representative transcripts in 10 of the MIKC_C_ clades, except for FLC-like, AGL15-like, DEF-like and StMADS11-like. Similar to previous reports, FLC-like and AGL15-like clades consist only *Arabidopsis* genes, suggesting eudicot specific lineages (Arora et al., 2007; Ballerini et al., 2013; Becker & Theißen, 2003; T. Zhao et al., 2006). Three groups consist *I. atropurpurea* sequences but lack *I. fulva* representatives, AGL12-like, AGL17-like, and GMM13-like. AGL17-like and GMM13-like are not supported by the bootstrap analysis. AGL12-like has three *I. atropurpurea* transcripts, and this clade was well supported. AGL12-like and AGL17-like genes are involved in root development (Tapia-López et al., 2008; H. Zhang & Forde, 1998), and while the *I. atropurpurea* sequences were derived also from root tissue, the *I. fulva* transcriptome was based on floral and leaves tissues (Ballerini et al., 2013). Four *I. atropurpurea* sequences were clustered alone in a well supported group (81%, designated “Unknown”). These sequences might be of genes unique to *I. atropurpurea.*

Within most of the clades *I. atropurpurea*, *I. fulva* and rice grouped together and *Arabidopsis* sequences grouped together, suggesting a strong species and monocot/eudicot homology. In Arora et.al. *Arabidopsis* and rice also cluster together within the type I MADS clades (Arora et al., 2007). Furthermore, a phylogeny of representative type I and II MADS-box genes from several distantly related plant species also showed similar monocot/eudicot separation within clades (Gramzow & Theissen, 2010).

### R2R3-MYB genes

We found 256 transcripts in the *I. atropurpurea* transcriptome that were annotated as belonging to the MYB family by either trinity or PFAM. Sixty-seven of them were found to have the R2R3-MYB domain. The rest of the transcripts most likely belong to other MYB groups such as R1-MYB, MYB-like proteins, etc, and some might also be incomplete sequences.

To create the phylogenetic tree, we aligned the transcripts against R2R3-MYB sequences from *Arabidopsis,* rice, and *I. Fulva* (figure 6). R3-MYB (R1R2R3) is another major MYB type, that was either the origin of R2R3-MYBs in plants (Rosinski & Atchley, 1998) or evolved from R2R3-MYB (Jiang, Gu, Chopra, Gu, & Peterson, 2004), and was also included in the phylogenetic analysis. To analyze the tree, we mainly followed the classification made by Ballerini et al., which consist *Iris* sequences (Ballerini et al., 2013). The organization of the clades in the dendrogram corresponds with that in Ballerini et al., with 26 of the groups supported by bootstrap (>50%). Several groups showed differences from the phylogeny in Ballerini et al., mostly in the form of a sequence clustered to a different clade, and in most cases not supported by bootstrap. Some major differences were observed for example in group 10, which was separated into 2 clades in our analysis, one with the *Arabidopsis* sequences and one with rice. Similar separation was found in Du et.al., in which the rice sequences are in a separated clade with *Zea Maize*, designated as S42 (Du et al., 2015). In our tree, group 16 was also separated into 2 clades, in accordance with other published MYB trees (Du et al., 2015; Yanhui et al., 2006).

**Figure 6.**
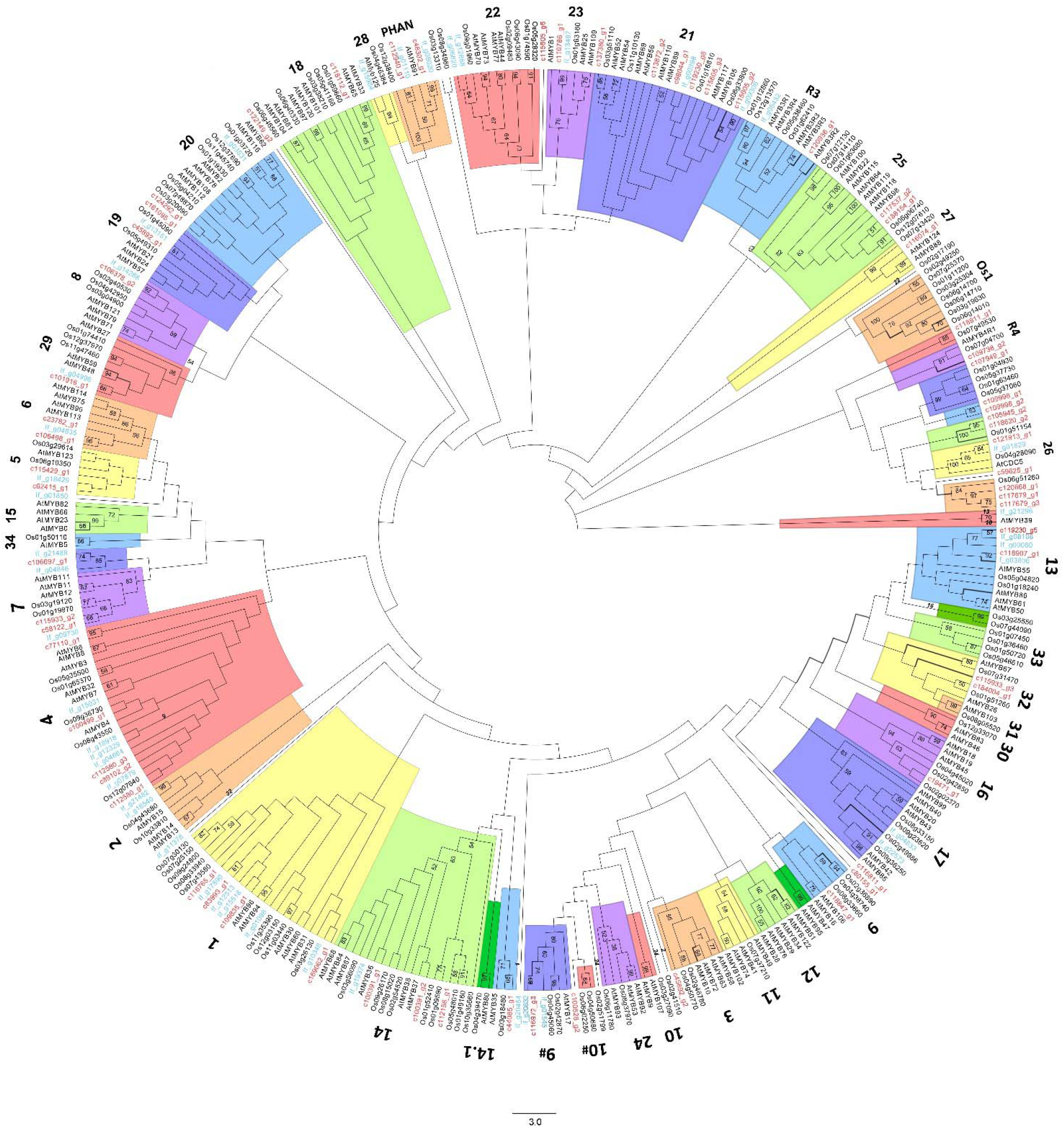
Phylogenetic analysis of R2R3-MYB proteins from the *Iris* transcriptome (highlighted in red), *I. fulva (If), Arabidopsis,* encoded by *AtMYB*, and rice (Oryza sativa, Os). *I. atropurpurea* transcripts names are in red. Colours are for visual separation only.

Fourteen groups of R2R3-MYB genes in the phylogenetic tree lack *I. atropurpurea* representatives, whereas in nine of them *I. fulva* representatives were also lacking, suggesting gene lineages that might not exist in Irises (11, 12, 15, 24, 30, 31, 33, 34, and Os1). Consistent with previous phylogenetic studies, groups 12 and 15 also lack rice representatives, suggesting eudicot specific lineages (Ballerini et al., 2013) (Yanhui et al., 2006). A comparative analysis of R2R3-MYBs from 50 major eukaryotic lineages showed that group 12 consists only of *Arabidopsis* sequences and that group 15 consists only of eudicot species (Du et al., 2015). Genes in these groups have been shown to control trichome initiation in shoots, root hair patterning, and Cruciferae-specific glucosinolate biosynthesis (Ambawat et al., 2013; Dubos et al., 2010; Y. Li et al., 2013). Two of the groups lacking representatives from *Iris*, 33 and Os1, consist only rice genes. Genes from group 33 were previously designated in a monocot-specific clade together with corn (*Zea maize*) sequences (Du et al., 2015). Several groups had only *I. atropurpurea* representatives, lacking *I. fulva,* and vice versa. In addition, in contrast with our expectations, only in a few of groups *I. atropurpurea* and *I. fulva* clustered together within the clade. These observations further support the phylogenetic distance between the two species.

We found two new (bootstrap supported) subgroups consisting only rice and *I. atropurpurea* sequences. Previous phylogenetic studies in other plant species also identified new R2R3-MYB subgroups with no *A. thaliana* representatives. These subgroups might represent genes with specialized functions which were either lost in *Arabidopsis* or obtained after the divergence from the last common ancestor (Ballerini et al., 2013; Wilkins, Nahal, Foong, Provart, & Campbell, 2009). Several *I. atropurpurea* sequences did not cluster together with R2R3-MYBs from any other species, including *I. fulva*. This suggests that these MYB genes might have been acquired in *I. atropurpurea* after divergence within the *Iris* group.

Other MYB and MADS gene groups, which were not identified in our transcriptome, could be genes that were not conserved in irises. Alternatively, these genes might be expressed in earlier stages of flowering initiation, before the appearance of buds (Perl, 1984), and thus undetected in the transcriptome. In *Iris lortetii*, it was shown that flower organs genes are mostly expressed in an early stage, about two months before stem elongation, when the flower meristem is hidden in the rhizome (Perl, 1984). Possibly this is the stage when more flower development genes can be found; however, this stage was not sampled in this study and will be explored in further research.

### Development and characterization of cDNA-derived SSR markers

For the development of new molecular markers, we used all of the 258,466 contigs, generated in this study, to mine potential microsatellites. We defined microsatellites as di- to hexanucleotide SSR with a minimum of four repetitions for all motifs. We identified 1,503 potential SSRs in 1,241 contigs, of which 263 sequences contained more than one SSR. Only 164 of the contigs containing SSRs had annotation and were annotated to 115 genes. We assessed the frequency, type, and distribution of the potential SSRs (figure 7). The SSRs included 924 (61.5%) di-nucleotide motifs, 396 (26.4%) tri-nucleotide motifs, 173 (11.5%) tetra-nucleotide motifs, 10 (0.7%) penta-nucleotide motifs, and zero (0%) hexa-nucleotide motifs. The di-, tri-, tetra- and penta-nucleotide repeats had 8, 30, 37 and 9 types of motifs, respectively. The most abundant di-nucleotide type was GA/TC (254, 16.9%), followed by AG/CT (197, 13.1%) and AT/AT (159, 10.6%). The most abundant tri-nucleotide repeat type was TTC/GAA (37, 2.5%).

**Figure 7.**
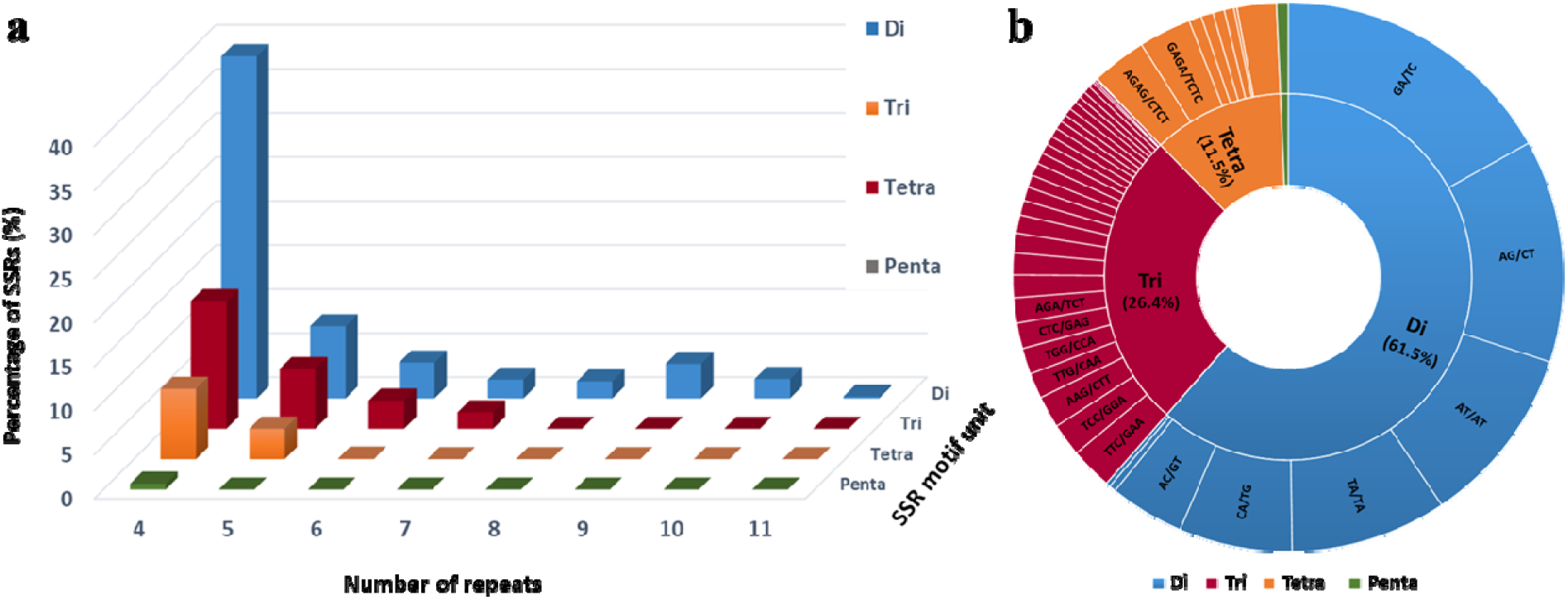
Characterization of SSRs loci found in *Iris* transcriptome. a. Distribution of SSR motif repeat numbers and relative frequency. b. Frequency distribution of SSRs based on motif sequence types.

Di-nucleotide SSRs are usually more common in genomic sequences, whereas tri-nucleotide SSRs are more common in RNA sequences (Luo et al., 2005; Thiel, Michalek, Varshney, & Graner, 2003; Varshney, Graner, & Sorrells, 2005; Varshney, Thiel, Stein, Langridge, & Graner, 2002). Also, tri-nucleotide repeats are more abundant than dinucleotide repeats in plants (Varshney et al., 2005). However, in our SSRs, the di-nucleotide repeat type was the most abundant motif detected of all repeat lengths. A higher number of di-nucleotide repeats in RNA sequences has been reported in Louisiana irises (Tang et al., 2009), and in other plants such as rubber trees (D. Li, Deng, Qin, Liu, & Men, 2012) and *Cajanus cajan* (pigeonpea) (Raju et al., 2010). The most abundant di- and tri-nucleotide motifs in *I. atropurpurea* were GA/TC and TTC/GAA, respectively. These results were also coincident with SSRs developed for Louisiana irises, in which the most abundant di- and tri-nucleotide motifs are AG/CT and AAG/CTT (Tang et al., 2009).

Until now, SSRs in irises were reported only for Louisiana and Japanese irises (Sun et al., 2012; Tang et al., 2009); however, these SSRs were not transferable to *Oncocyclus* irises (Y. Sapir, un-published). The relatively large set of SSRs obtained from the *I. atropurpurea* transcriptome may enable development of markers for population genetic studies in the Royal Irises.

## Conclusions

In this study, we reported a comprehensive characterization of the transcriptome of *Iris atropurpurea*, an important emerging model for understanding evolutionary processes (Arafeh et al., 2002; Dorman et al., 2009; Lavi & Sapir, 2015; Sapir & Mazzucco, 2012; Sapir et al., 2005, 2006; Volis et al., 2010; Carol A. Wilson et al., 2016; Yardeni et al., 2016). Although transcriptome based on a single replication cannot enable gene expression analysis and extensive biological conclusions, the *Iris* transcriptome established in this study provides a useful database that will increase the molecular resources for the Royal Irises. These resources are currently available only for other iris species (Sun et al., 2012; Tang et al., 2009), which despite belonging to the same genus, they are quite distant from the Royal Irises, hence not easily transferable. In the past decade, many studies have been using transcriptome *de novo* sequencing and assembly to generate a fundamental source of data for biological research (Ballerini et al., 2013; Kamenetsky et al., 2015; Meyer et al., 2009; S. Tian et al., 2015; J. Zhang et al., 2012). We generated a substantial number of transcript sequences that can be used for the discovery of novel genes, and specifically genes involved in flower development in irises.

While we did not perform a complete analysis of MADS and R2R3 MYB evolution, we mainly aimed to identify flower development genes and classify their function, and thus provide a framework for the *Iris* genes sequenced in this study. The numerous SSR markers identified will enable the construction of genetic maps and answering important questions in population genetics and conservation. Although genetic studies are still in their early stages in the Royal Irises, we believe that our transcriptome will significantly support and encourage future evolutionary-genetic research in this ecologically important group.

## Acknowledgments

We thank the Technion Genome Center for technical assistance and the Bioinformatics Core Facility at Ben-Gurion University for their assistance in the bioinformatics analysis. We thank N. Kane for the Perl script for mining SSRs from transcriptome sequences. This research was funded by the Israel Science Foundation to YS (grant No. 336/16).

## Compliance with Ethical Standards

### Funding

This research was funded by the Israel Science Foundation (grant No. 336/16). Competing interests: The authors declare that they have no conflict of interests.

## Authors’ contributions

YBL and YS designed the study and drafted the manuscript. YBL conducted the experimental work. YBL, ES, and MPS carried out the bioinformatics analysis. All authors read and approved the final manuscript.

## Data availability

All data generated or analyzed during this study are available at NCBI’s Gene Expression Omnibus (GEO), accession number GSE121786.

